# Spatial organization of cells and variable expression of autophagy, apoptosis, and neurodevelopmental genes might underlie selective brain region vulnerability in Attention-Deficit/Hyperactivity Disorder

**DOI:** 10.1101/652792

**Authors:** Jonathan L. Hess, Nevena V. Radonjić, Jameson Patak, Stephen J. Glatt, Stephen V. Faraone

**Affiliations:** Department of Psychiatry, SUNY Upstate Medical University; Syracuse, NY, USA; Department of Neuroscience, SUNY Upstate Medical University; Syracuse, NY, USA

**Keywords:** ADHD, autophagy, apoptosis, reactive oxygen, neurodevelopment, brain volumes, cortical thickness, imaging genomics, microarray

## Abstract

Genetically influenced changes in brain organization occur over the course of development. Large-scale brain imaging studies by the ENIGMA Consortium identified structural changes associated with attention-deficit/hyperactivity disorder (ADHD). It is not clear why some brain regions are impaired and others spared by the etiological risks for ADHD. We hypothesized that spatial variation in brain cell organization and/or pathway expression levels contribute to selective brain region vulnerability (SBRV) in ADHD. In this study, we used the largest available collection of imaging results from the ADHD ENIGMA Consortium along with high-resolution *postmortem* brain microarray data from Allen Brain Atlas from 22 sub-cortical and cortical brain regions to investigate our selective brain region vulnerability (SBRV) hypothesis. We performed deconvolution of the bulk transcriptomic data to determine abundances of neuronal and non-neuronal cells in the brain. We then assessed the relationships between gene set expression levels, cell abundance, and standardized effect sizes representing regional changes in brain sizes in cases of ADHD. Our analysis yielded significant correlations between apoptosis, autophagy, and neurodevelopment genes with reductions in regional brain sizes in ADHD, along with associations to regional abundances of dopaminergic neurons, astrocytes, oligodendrocytes, and neural progenitor cells. This works provides novel mechanistic insights into SBRV in ADHD.

## Introduction

Attention-deficit/hyperactivity disorder (ADHD) is a highly heritable neurodevelopmental disorder characterized by high levels of inattention, impulsivity, and hyperactivity which frequently persist into adulthood (1). ADHD is a relatively common disorder that affects about 5% of school-age children and 2.5% of adults. Many genetic and environmental risk factors for ADHD have been documented but the mechanisms ultimately leading to the symptoms of the disorder are unknown (1).

Multi-site structural magnetic resonance imaging (**sMRI**) mega-analyses performed by the ENIGMA Consortium identified sub-cortical regions with significant smaller volumes in children diagnosed with ADHD compared to unaffected comparison participants, including: the accumbens, amygdala, caudate, hippocampus, and putamen. Stratification by age group revealed that sub-cortical volumetric reductions were more prominent for children with ADHD than for adults with the disorder (2). More recently, the ENIGMA Consortium identified reductions in cortical thickness within the fusiform, precentral and paracentral gyri, entorhinal cortex, and parahippocampal lobe in cases of ADHD spanning multiple age groups. Stratifying individuals into different age groups showed that children with ADHD show substantially greater reductions in cortical thickness compared to adults with ADHD. These findings reported by the ENIGMA Consortium (2) join a growing body of work that finds that ADHD-associated deficits in regional brain volumes and cortical thickness attenuate with aging (3, 4).

Our hypotheses about selective brain region vulnerability (**SBRV**) derive, ultimately, from the theory of pathoclisis introduced in 1922 (5), which posited that the basis of selective vulnerability to disease may be explained by variation in cell types and biochemical pathways of brain regions. We postulate that mechanisms underlying SBRV explain why some brain regions show structural brain changes associated with ADHD while others do not (2, 6). Previously, we found that autophagy, apoptosis, and oxidative stress pathways were associated with smaller sub-cortical brain region volumes in ADHD and suggested that they may mediate SBRV (7). The present work builds on our previous study by including newly released data from the ENIGMA-ADHD working group for cortical brain regions. We also investigate the brain cell types that may mediate SBRV in ADHD. Our goal was to determine if a distinct gene expression or cellular profile was characteristic of brain regions implicated in ADHD. Such data could shed light on etiological mechanisms underlying SBRV.

## Methods

A diagram summarizing our analytic framework is provided in **Figure 1.**

**Figure 1.**
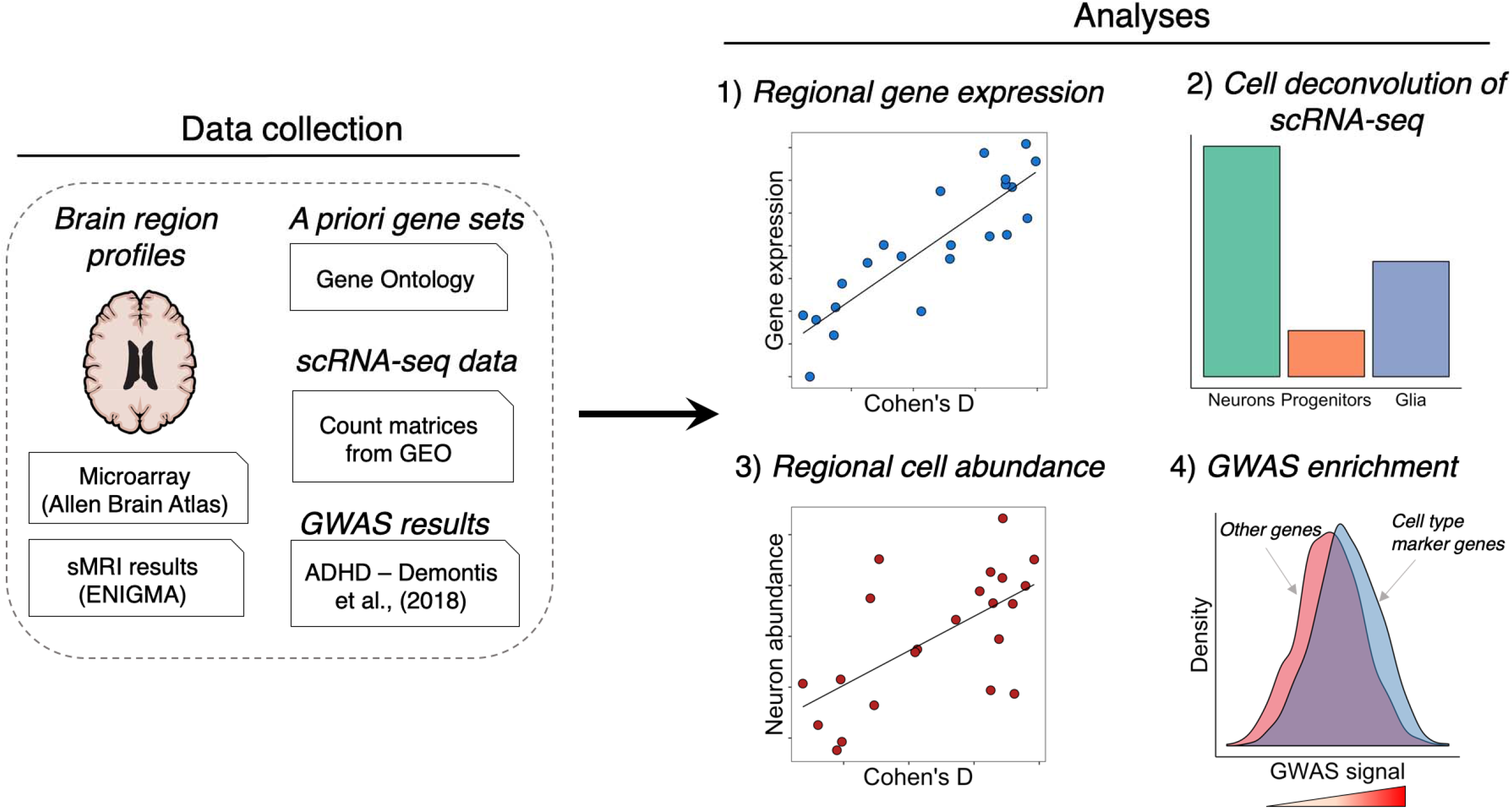
A diagram showing the general workflow used for our study. We collected regional *postmortem* brain transcriptome profiles from Allen Brain Atlas and summary statistics from two neuroimaging meta-analyses by the ENIGMA Consortium for ADHD. We analyzed the *postmortem* brain data for correlations between gene set expression levels of apoptosis, autophagy, neurodevelopment, neurotransmitter, oxidative stress and ADHD candidate genes with volumetric reductions seen in ADHD cases from the ENGIMA studies. We also deconvoluted the regional *postmortem* brain transcriptome profiles into discrete cell types and tested for a correlate between cell abundance and volumetric reductions in ADHD. We tested for enrichment of GWAS signals associated with ADHD among gene sets of interest (i.e., cell type marker genes, Gene Ontology gene sets). **Abbreviations**: Gene Expression Mmnibus (GEO), genome-wide association study (GWAS), single cell RNA-sequencing (scRNA-seq).

### Gene sets

The latest version of the Gene Ontology (GO) database was downloaded *via R* package *GO.db* (version 3.7.0) (8) to retrieve the most up-to-date annotations for the following gene sets of interest that had previously been implicated in ADHD and that we hypothesized might be involved in SBRV: apoptosis, autophagy, neurotransmission, neurodevelopment, and reactive oxygen species (9–12). A total of 1,313 unique HGNC genes were identified including 43 ADHD candidate genes designated in our previous study (7). 1,288 of these had expression data in the Allen Brain Atlas. **Supplementary Figure 1** shows the number of genes contained in each gene set and the amount of pairwise overlap between gene sets.

### Structural neuroimaging data for cortical and subcortical brain regions

We used data from two international multi-site neuroimaging meta-analyses conducted by the ENIGMA-ADHD working group that evaluated structural T1-weighted brain MRI data from individuals diagnosed with ADHD and unaffected comparison participants. Summary statistics denoting case-control differences in cortical thickness and sub-cortical volume as standardized effect sizes (Cohen’s *D*) were obtained from published papers by the ENIGMA Consortium (2, 6). Structural MRI measures had been generated using a validated MRI processing protocol with the software *FreeSurfer*. Differences in cortical thickness and subcortical volumes found between cases and unaffected comparison participants had been adjusted for age, sex, and total intracranial volume.

### Postmortem brain transcriptome profiles from adult donors

Pre-processed microarray data was downloaded from the Allen Brain Atlas (13) containing the transcriptome profiles for 22 brain regions that were evaluated by the ENIGMA-ADHD working group (15 cortical and 7 sub-cortical). A total of 58,692 probes were used to assay transcriptome expression profiles from neuropathologically normal *postmortem* brain specimens harvested from six adult donors (13). Probe-level data were collapsed down to 19,274 HGNC genes by calculating the median expression level of probe clusters (*i.e.*, set of probes that measure a single gene) within each donor and brain region. Expression levels were then summarized across gene sets by computing the mean expression levels of genes within each gene set, which followed the same technique from our previous study (7). A plot of normalized expression values across regions of interest from Allen Brain Atlas is provided in **Supplementary Figure 2**.

### Developmental transcriptomes

Normalized RNA-sequencing data generated from *postmortem* brain tissues was downloaded from the web portal of the Developmental Transcriptome of the Allen Brain Atlas (14), providing region-specific transcriptome-wide expression levels within 21 cortical and 5 sub-cortical brain structures in prenatal (8 to 37 post-conceptual weeks) and postnatal donors (4 months to 40 years). Average expression levels were calculated for the six gene sets separately across cortical and sub-cortical regions within each age group.

### Deconvolution of bulk brain tissue transcriptomes into specific brain cell types

An analytic approach called “cell deconvolution” was performed to infer the relative abundance of neuronal and non-neuronal cell fractions from bulk transcriptome data obtained from the Allen Brain Atlas. This procedure required single-cell RNA-sequencing (**scRNA-seq**) data from independent samples to determine the parameters for cell deconvolution. Read count matrices were downloaded from the Gene Expression Omnibus (**GEO**) for two scRNA-seq studies of major neuronal, glial, and vascular cells in the human brain; one of the studies performed transcriptome sequencing of ventral midbrain cells harvested from human embryos (GEO accession ID: GSE76381) (15), while the other sequenced the transcriptome of brain cells derived from the adult cortex (GEO accession ID: GSE67835) (16). Standard quality control procedures were applied to the raw read count data using the software *edgeR* (17) to minimize technical variation that could obscure biological signals, including: discarding genes that were expressed near background levels (≤1 read counts in > 90% of the sample), and transforming raw read counts to the log_2_ counts per million (**CPM**) scale and performing quantile normalization. We used the software *dtangle* (18) to identify cell-type marker genes (*i.e.*, genes expressed above the 90^th^ percentile) and perform deconvolution on expression data from each of the 22 brain regions to estimate abundance of neuronal and non-neuronal cells.

### Statistical analysis

Pearson’s correlation tests were used to determine the association between Cohen’s *D* values that reflect brain morphometric changes in ADHD cases, separated by the four case-control age groups evaluated by the ENIGMA Consortium and gene set expression levels in *postmortem* sub-cortical and cortical brain regions obtained from the Allen Brain Atlas. The Nyholt procedure was used to correct for multiple comparisons (19), which accounts for the correlation structure between variables, thus is considered beneficial for statistical power. We applied the Nyholt procedure to the correlation matrix of gene set expression levels and corrected for the number of multiple comparisons performed within each case-control age group defined by the ENIGMA-ADHD working group. A *post hoc* correlation test was performed to examine the relationship between Cohen’s *D* effect sizes and expression levels on a per-gene basis, wherein the Bonferroni procedure was used, which is relatively more conservative compared to the Nyholt method, in order to uncover genes with the strongest association with brain morphometric changes in the brains of ADHD cases among those gene sets already associated with ADHD-related brain changes (*i.e.*, Bonferroni *p* < 0.05). Pearson’s correlation test was used to examine the relationship between abundance of specific brain cell types across sub-cortical and cortical brain regions along with gene set expression levels, wherein the effective number of independent tests was the dot product of the number of independent cell types and the number of gene sets evaluated. Lastly, Pearson’s correlation tests were performed on abundance of specific brain cell types and Cohen’s *D* effect sizes for brain structural changes in ADHD within each of the case-control age groups. The Nyholt procedure was used to compute the effective number of independent cell types then adjust the significance threshold to account for multiple testing. A *post hoc* multivariate regression model tested whether multiple significant gene sets had conditionally independent associations with Cohen’s *D* values.

### Gene set analysis with GWAS data for ADHD

We performed a gene-level and gene set association analysis using the largest available GWAS meta-analysis results for ADHD (20). Identifiers for over 3 million single nucleotide polymorphism (**SNPs**; rsIDs) and *p*-values were supplied to *MAGMA* (v1.07) (21) to perform quality control of summary statistics and compute *z*-scores for gene-level associations with ADHD by averaging the observed significance values for intragenic SNPs after adding the same window definitions of a gene (−10 kilobases upstream of the transcription start site and +35 kilobases downstream of the transcription stop position) used by the Psychiatric Genomics Consortium (22). Gene-level associations were computed for a total of 20,274 genes, of which 1,171 were included in our gene sets of interest. A genome-wide significance threshold was set at 2.47×10^−6^ for the gene-level tests (*i.e.*, 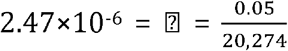). With *MAGMA*, a linear regression model was used to perform a competitive gene set enrichment analysis testing for differences in gene-level association scores for ADHD between a gene set and all other genes in the genome while covarying for minor allele count, gene length, number of SNPs per kilo base of the gene, and linkage disequilibrium between genes.

## Results

### Gene sets associated with reduced brain sizes in ADHD

Similar patterns of developmental expression levels were found within each gene set across cortical and sub-cortical brain structures (**Supplementary Figure 2**). Expression levels of autophagy, neurodevelopment, neurotransmission, and oxidative stress gene sets increased over the lifespan for both cortical and sub-cortical structures (**Supplementary Figure 2**). ADHD candidate genes displayed a prominent increase in expression following birth that steadily increased into adulthood. Conversely, apoptosis genes showed a gradually decreasing expression beginning prenatally and extending into adulthood, with a period of up-regulation between 2 to 3 years of age.

Figure 2 shows the correlations between the Cohen’s *D* values indexing differences between ADHD and unaffected comparison (UC) participants in regional brain volumes and the gene set expression levels for the six *a priori* selected gene sets. As the summary statistics for the correlation tests in **Table 1** show, the expression levels of three gene sets were significantly negatively correlated with ADHD-associated volumetric changes: autophagy (children with ADHD, Pearson’s *r* = −0.67, *p* = 7.0×10^−4^; all ages, Pearson’s *r* = −0.72, *p* = 2.0×10^−4^), apoptosis (children with ADHD, Pearson’s *r* = −0.53, *p* = 0.01; all ages, Pearson’s *r* = −0.66, *p* = 9.0×10^−4^), and neurodevelopment (adults with ADHD, Pearson’s *r* = −0.59, *p* = 4.0×10^−3^; all ages, Pearson’s *r* = −0.61, *p* = 2.4×10^−3^). The associations between the autophagy and apoptosis gene sets with ADHD-associated volumetric changes disappeared when the two gene sets were both specified as predictors in a *post hoc* multiple linear regression model that used Cohen’s *D* values from ENGIMA’s analysis of all ADHD cases and unaffected comparison participants as the outcome variable (*p*-values = 0.072 and 0.76, respectively), suggesting that these gene sets are not contributing conditionally independent effects on ADHD-associated volumetric changes. This finding is consistent with the fact that the autophagy and apoptosis gene sets share a significantly greater number of genes than expected by chance (**Supplementary Figure 1**). Oxidative stress genes were nominally significantly associated with ADHD-associated volumetric changes prior to multiple testing correction in the child, adolescent, and combined sample of ADHD cases and unaffected comparison participants (uncorrected *P*_*S*_ = 0.02, 0.02, and 0.03, respectively).

**Figure 2.**
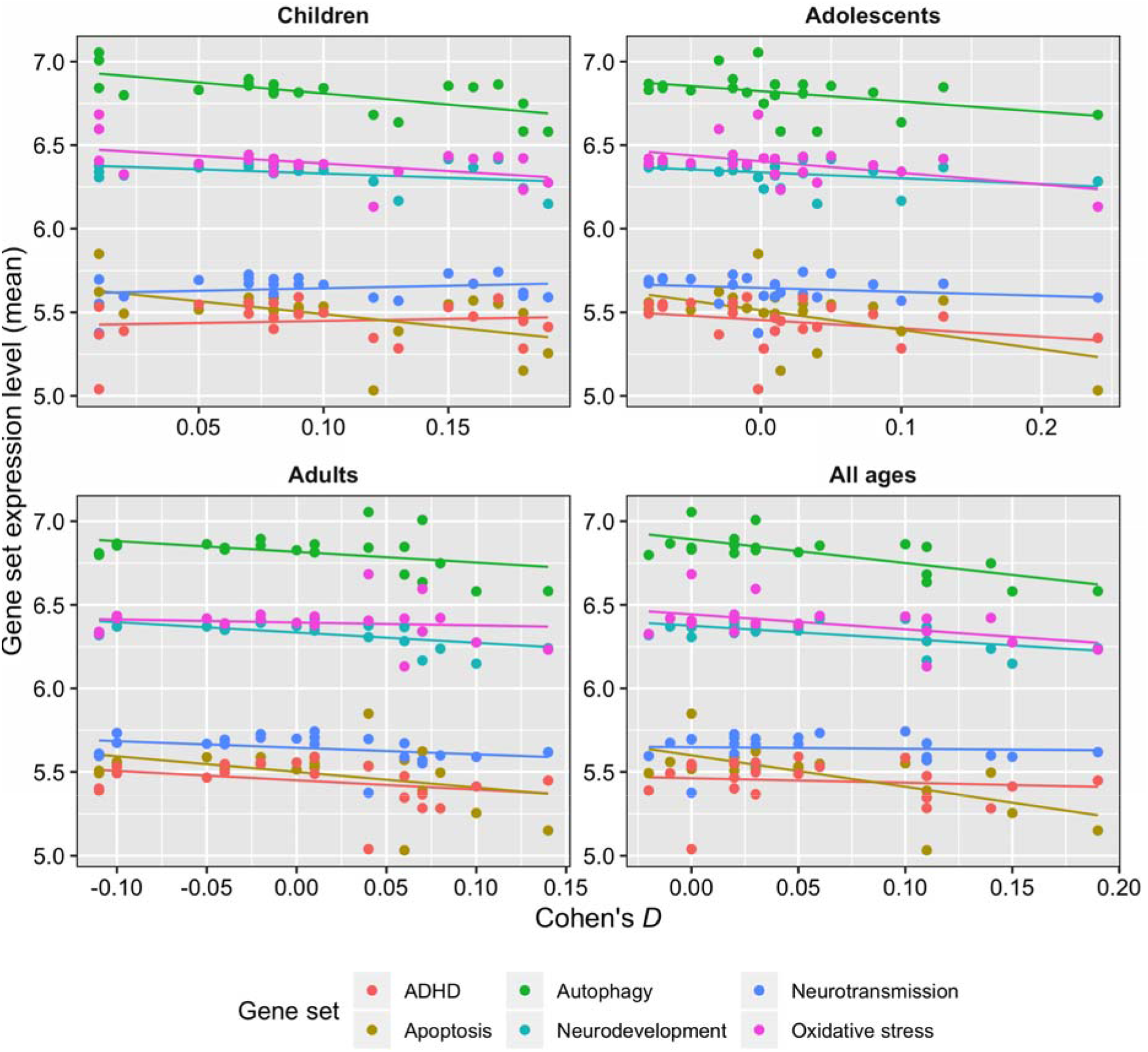
Scatterplots and best-fit regression lines show correlations between volumetric differences in ADHD cases across 22 brain regions (x-axis, Cohen’s *D*) with gene set expression levels (y-axis). Cohen’s *D* effect sizes were stratified according to the age group described by the ENIGMA-ADHD working group in their case-control analyses of sub-cortical and cortical brain MRI data. Dots are color coded by gene set. Table 1 gives the statistical significance of these correlations.

**Table 1.**
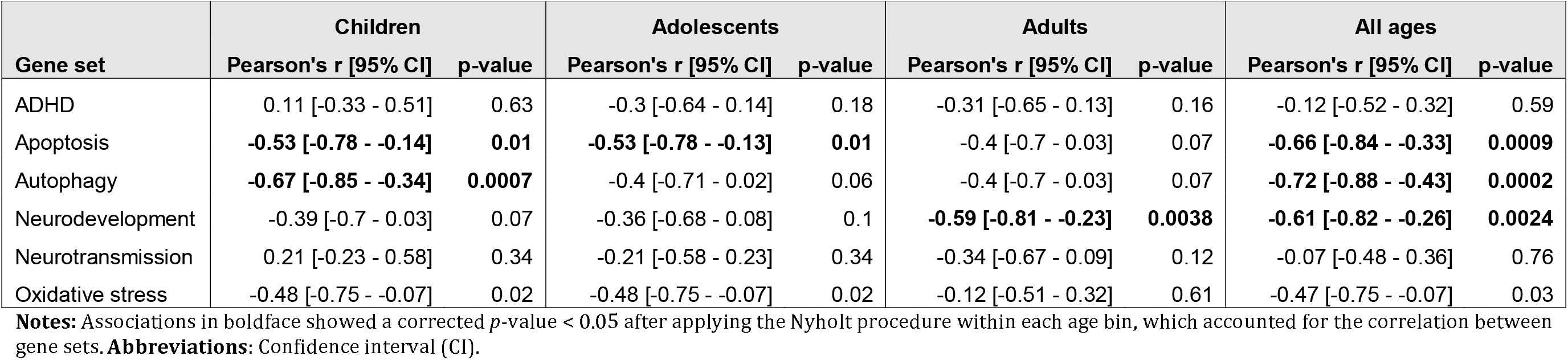
Correlation of gene set expression levels with volumetric changes in ADHD cases across 22 brain regions.

The distributions of Pearson’s correlation coefficients describing the relationship between expression levels of 974 genes from the autophagy, apoptosis, neurodevelopment, and oxidative stress gene sets and ADHD-associated volumetric changes are in **Supplementary Figure 3**. Sixty-four genes were significantly correlated with Cohen’s *D* effect sizes at a Bonferroni *p* < 0.05 with the following number of genes associated brain volumetric changes in ADHD cases for the following age groups: 2 for children, 1 for adolescents, 15 for adults, and 46 for all ages combined (**Supplementary Figure 4**). Expression levels of *TAOK2*, a serine/threonine kinase gene, had the most significant association with ADHD-associated volumetric reductions (Pearson’s *r* = −0.85, uncorrected *p* = 6.13×10^−7^, Bonferroni *p* = 6.0×10^−4^).

### Association of cell type abundance with gene sets and volumetric changes in ADHD

**Figure 3** shows significant associations found between 14 brain cell types and autophagy, apoptosis, neurodevelopment, or oxidative stress genes (Nyholt *p* < 0.05). The cell types implicated by these associations were: four glial cell types (astrocytes, embryonic microglia, and embryonic oligodendrocytes), microvasculature cells (embryonic pericytes), five neuron types (embryonic dopaminergic neurons, embryonic red nucleus neurons, embryonic GABAergic neurons, embryonic serotonergic neurons, and adult neurons), and four progenitor cell types (embryonic radial glial cells, embryonic lateral floorplate progenitor cells, embryonic midline progenitor cells, embryonic mediolateral neuroblasts, and adult oligodendrocyte progenitor cells).

**Figure 3.**
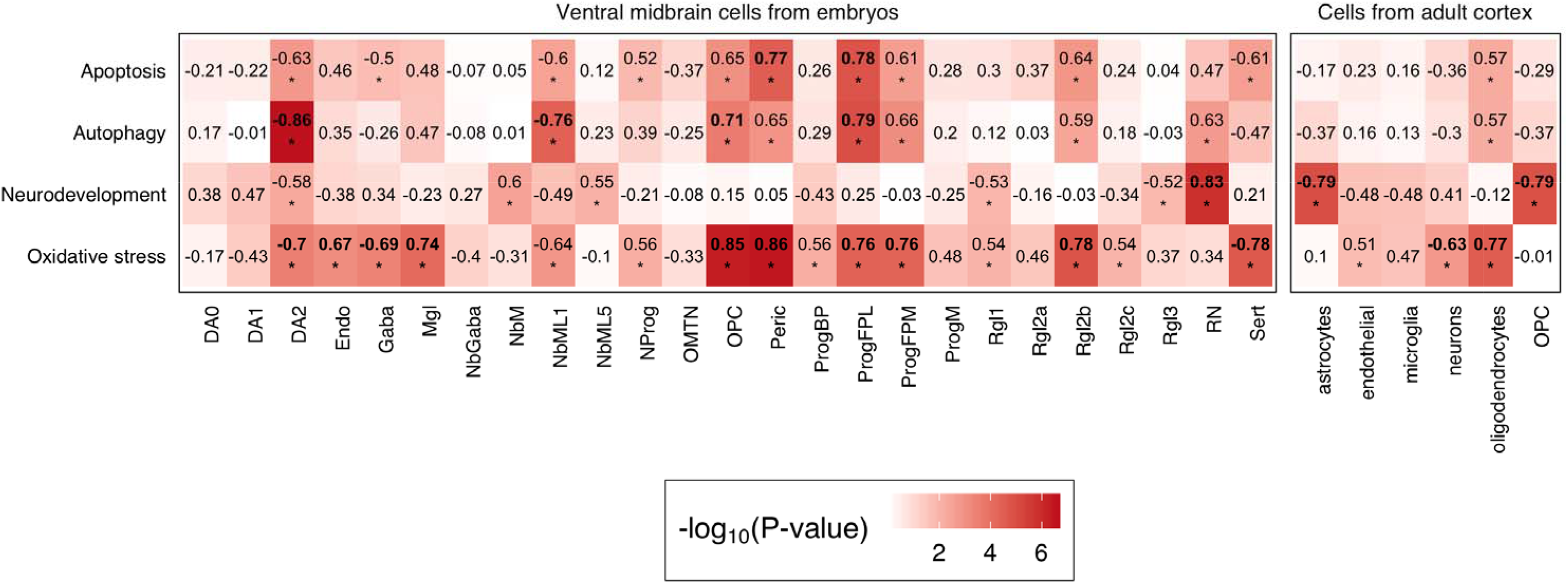
Correlation of gene set expression levels with estimate abundances of neuronal and non-neuronal cells identified from scRNA-seq studies of embryonic ventral midbrain specimens (left panel) and adult cortex (right cortex). Associations marked by asterisks (*) reached a significance level of *p* < 0.05, whereas associations appearin in boldface were statistically significant after multiple testing correction (Nyholt *p* < 0.05). **Abbreviations**: dopaminergic cells (DA0-2), endothelial cells (endo), GABAergic neurons (Gaba), microglia (Mgl), mediolateral neuroblasts (NbML1-5), neural progenitor (NProg), oculomotor and trochlear nucleus (OMTN), oligodendrocyte progenitor cells (OPC)progenitor (Prog), pericytes (Peric), progenitor medial floorplate (FPM), midline (M), basal plate (BP), radial glial-like cells (Rgl1-3), red nucleus (RN), serotonergic cells (Sert).

Significant associations were found between five embryonic and two adult brain cell types and ADHD-associated volumetric changes (Nyholt *p* < 0.05, **Figures 4** – **5**). The cell types implicated by these associations were: embryonic lateral floorplate progenitor cells, embryonic radial glial-like cells, embryonic dopaminergic neurons, embryonic GABAergic neuroblasts, embryonic midline neuroblasts, adult astrocytes, and adult oligodendrocyte progenitor cells.

**Figure 4.**
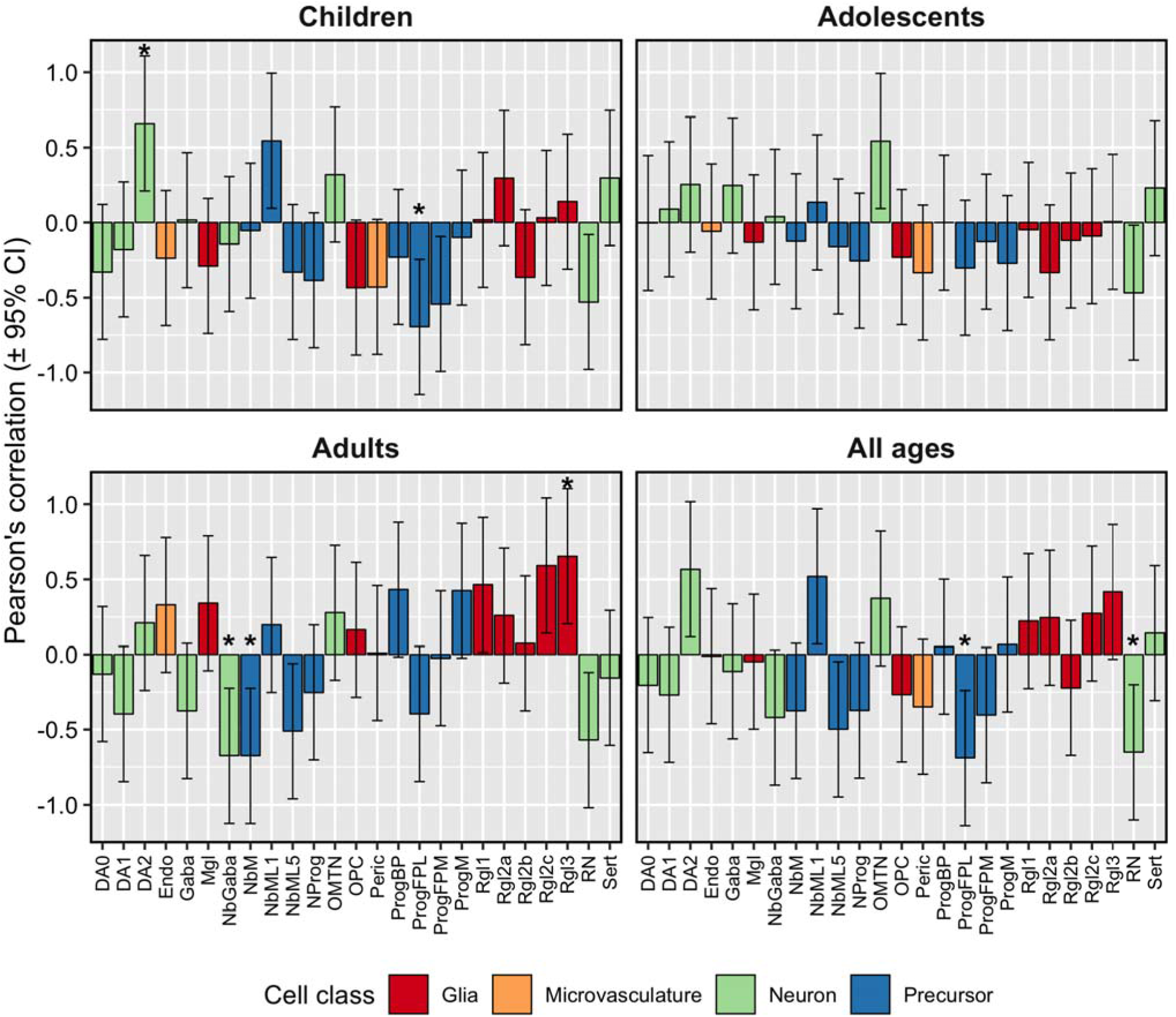
Estimates of neuronal and non-neuronal cell abundance based on deconvolution of Allen Brain Atlas guided by scRNA-seq from embryonic ventral midbrain specimens show correlations with volumetric differences in brain regions between ADHD cases and unaffected comparison participants. The asterisks (*) denote correlations that remained significant after correction for multiple testing with the Nyholt procedure applied within each age group. **Abbreviations**: dopaminergic cells (DA0-2), endothelial cells (endo), GABAergic neurons (Gaba), microglia (Mgl), mediolateral neuroblasts (NbML1-5), neura progenitor (NProg), oculomotor and trochlear nucleus (OMTN), oligodendrocyt progenitor cells (OPC)progenitor (Prog), pericytes (Peric), progenitor medial floorplat (FPM), midline (M), basal plate (BP), radial glial-like cells (Rgl1-3), red nucleus (RN), serotonergic cells (Sert).

**Figure 5.**
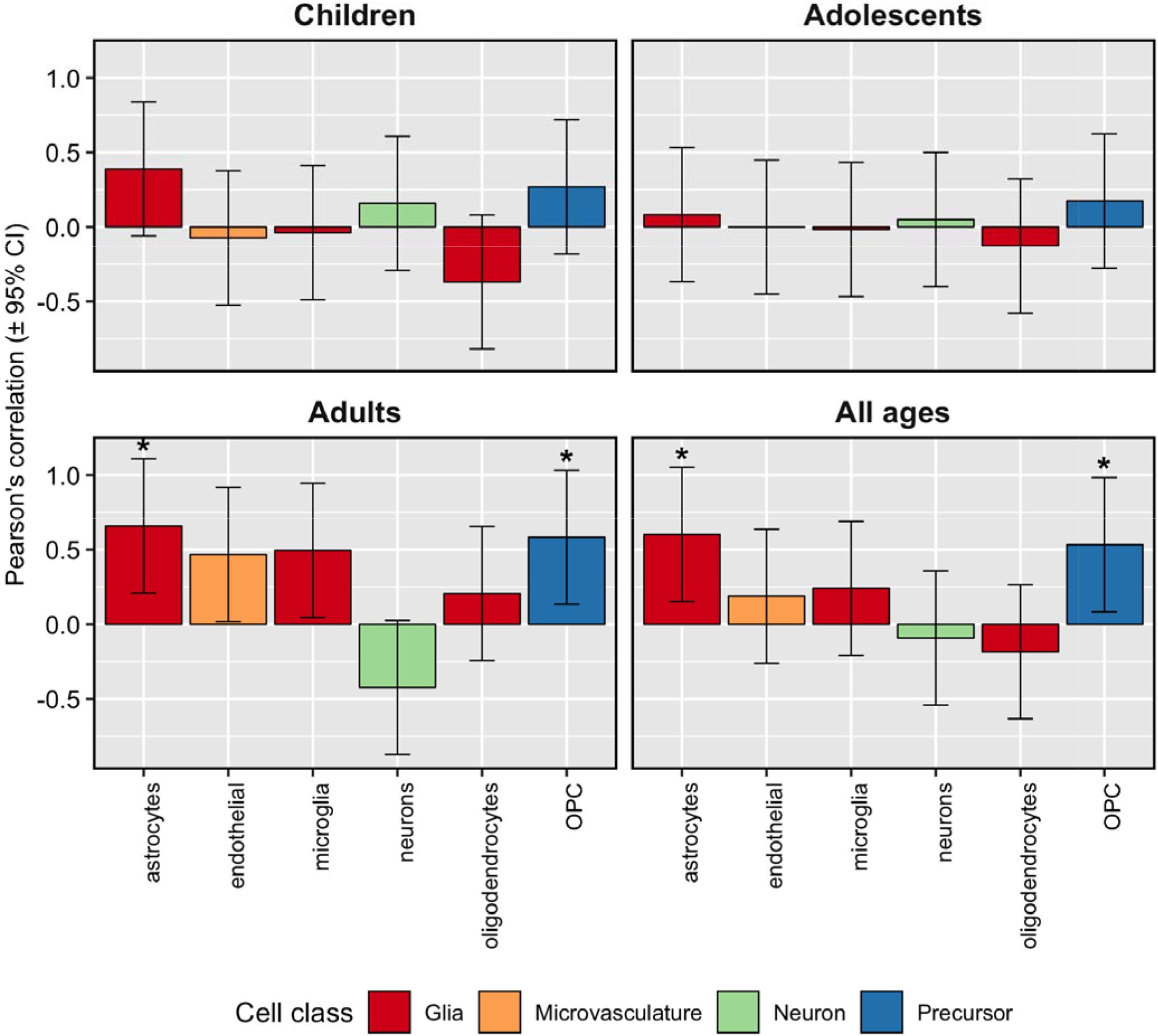
Estimates of neuronal and non-neuronal cell abundance based on deconvolution of Allen Brain Atlas guided by scRNA-seq data from adult cortex samples show correlations with volumetric differences in brain re ions between ADHD cases and unaffected comparison participants. The asterisks (*) denote correlations that remained significant after correction for multiple testing with the Nyholt procedure applied within each age group.

### Association of genes and gene sets with ADHD from GWAS meta-analysis

Genome-wide significant gene-level associations with ADHD were found for three genes, one of which belongs to the autophagy gene set (*KDM4A*, *p* = 2.61×10^−11^). The other two are members of the neurotransmission gene set (*CUBN*, *p* = 5.2×10^−7^ and *MEF2C*, *p* = 6.33×10^−7^). Competitive gene set enrichment did not uncover a significant association between autophagy, apoptosis, neurodevelopment, neurotransmission, or oxidative stress gene sets and ADHD (**Supplementary Table 1**). Marker genes for embryonic GABAergic neurons were significantly enriched with GWAS signals for ADHD (*p* = 0.029), but this association did not survive multiple testing correction (**Supplementary Table 1**).

## Discussion

This work sought to better understand selective brain region vulnerability (**SBRV**) in ADHD. We have extended our prior study of subcortical regions (Hess et al., 2018) by adding cortical data and by identifying the cell types possibly contributing to our results. Our results suggest that SBRV in ADHD may be due to temporo-spatial variation in the expression levels of autophagy and apoptosis genes. In adulthood, neurodevelopmental genes also appear to play a role in SBRV.

We previously reported a significant association between volumetric reductions in sub-cortical structures and spatial variation in expression of oxidative stress genes (7). However, after having nearly tripled our sample size, the association between oxidative stress genes and SBRV in ADHD was markedly diminished. Conversely, the significance of the associations between Cohen’s *D* and autophagy genes and apoptosis genes became stronger in this study compared to our previous findings (7), supporting the idea that the autophagy and apoptosis pathways (which share expressed genes) mediate SBRV for both cortical and sub-cortical structures. Although not parsimonious, it is plausible that neuronal or non-neuronal cells within cortical regions are more tolerant to oxidative stress compared to brain cells in sub-cortical regions and that oxidative stress my play a role only for sub-cortical structures.

Compared with our previous work, the association between SBRV in ADHD and the neurodevelopment gene set in adults is a novel finding. Neurodevelopmental genes play a critical role in the proliferation and differentiation of progenitor cells and the formation of neural circuits during fetal life and infancy. These genes, however, also contribute to neurogenesis later in life. The developmental expression data from BrainSpan show that neurodevelopmental genes are up-regulated during the early postnatal period, adolescence and in early adulthood (**Supplementary Figure 5**). 39 of the 263 genes (hypergeometric enrichment *p*-value = 2.5×10^−29^) in the neurodevelopmental gene set showed evidence of regulating adult neurogenesis based on the Mammalian Adult Neurogenesis Gene Ontology (**MANGO**) database (23). Taken together, we hypothesize that up-regulation of neurodevelopment genes during adulthood might promote neurogenesis to compensate for reduced brain volumes early in life. This offers a possible mechanism for the finding that all of the structural brain deficits found in ADHD youth attenuate in adulthood (2, 6, 24).

Evidence of hippocampal neurogenesis in adult humans was first shown by Erikson et al. (25), with subsequent support by others (26, 27). During human brain development an active area of neurogenesis is the subventricular zone where progenitor radial glia cells (which were implicated in our deconvolution analyses) play a crucial role by giving rise to neurons, astrocytes and oligodendrocytes (28) and remain as neural stem cells into adulthood (29, 30). Although there is a growing body of evidence for adult neurogenesis, there is an on-going debate about the interpretation of those findings.

Among the top 64 genes correlated with SBRV in ADHD, expression of the gene encoding TAO kinase 2 (*TAOK2*) had the strongest correlation with SBRV for adults with ADHD. Spatial expression profiles taken from the latest RNA-sequencing data from Genotype-Tissue Expression (**GTEx**) version 7 (31) show that *TAOK2* is widely expressed in the central nervous system, especially the cerebellum. *TAOK2* is part of the neurodevelopment gene set and has been shown to regulate brain size and neural connectivity, and is present in the 16p11.2 microdeletion region, a rare structural variant that has been associated with range of neurodevelopmental phenotypes including autism, ADHD, and intellectual disability (32–36). Deletion of *TAOK2* in mice was found to lead to deficits in dendritic growth, synaptic formation, cortical layering, and autism-associated phenotypes (36). We postulate that altered regulation of *TAOK2* affects brain development and ultimately results in reductions in the size of brain regions seen in ADHD.

According to our cell deconvolution analysis, brain regions showing greater ADHD associated volumetric reductions during development also showed elevated numbers of astrocyte- and oligodendrocyte- precursor cells in adulthood. These results imply that compensation for volumetric loss seen in adulthood could be attributed not only to increases in the number of neurons, but also to a compensatory response of glial cells. Indeed, animal models of ADHD have shown increased number of GFAP (astrocyte-specific marker) positive cells in the spontaneous hypertensive rat model (37) and astrocytosis in thalamus and cortex of Git1^−/−^ mice (38). Importantly, white matter abnormalities have been reported in ADHD (39). Oligodendrocytes are the main myelinating cells in the central nervous system (40). Myelination accounts for approximately 40% of human brain parenchyma and could be an important contributor to the differences in brain volume (41). Numerous reports link autophagy, cell survival pathways and upstream signaling cascades, such as PI3K-mTOR, in the regulation of myelination (42).

Regions with greater ADHD-associated volumetric reductions in adults displayed lower levels of neuronal progenitor cells types, including embryonic GABAergic neuroblasts. Indirect evidence linking GABAergic cells to ADHD has been shown in our study and by others (43–46). We found a nominally significant enrichment of GWAS signals for ADHD among genes that are highly expressed in adult GABAergic cells but did not find enrichment for genes that are highly expressed in embryonic GABAergic neuroblasts. Reduced concentration of extracellular GABA has been detected in children with ADHD, therefore abnormal functioning GABAergic cells may be relevant to ADHD (43). Furthermore, a genetic association was uncovered between GABAergic genes and ADHD symptoms (Naaijen et al., 2017). These data suggest that the role of these cell types in ADHD should be further studied.

Several findings suggest that SBRV in ADHD may have a primarily environmental etiology. We found that none of the biological pathways we identified as associated with SBRV were associated with ADHD in gene set analyses of a very large GWAS of ADHD. The 64 genes that were significantly correlated with SBRV in ADHD were not significant as a gene set in the largest available GWAS of ADHD (20). Consistent with the idea that SBRV may have a primarily environmental etiology, another study used linkage disequilibrium (LD) score regression to show that, although ADHD was genetically correlated with intracranial volume (**ICV**; *r*_*g*_ = −0.23, *p* = 1.5×10^−4^), it was not genetically correlated with sub-cortical brain structure volumes corrected for ICV (47). Those findings indicate little or no overlap between the common genetic variants that cause ADHD and those that regulate subcortical SBRV. In contrast to these findings, variation in cortical thickness of the fusiform gyrus and inferior parietal and lingual cortices was genetically correlated with ADHD based on LD score regression (48), suggesting that common risk variants could play a small role in cortical SBRV in ADHD. The literature on rare variants in ADHD is too sparse (49, 50) to assess what role they might play in the disorder.

Given these weak genetic findings, it seems reasonable to postulate that environmental risks and their interactions with genes may explain SBRV in ADHD. This led us to propose the following theoretical model of SBRV in ADHD. Our model attributes the initial etiology of SBRV to environmental events that threaten neural integrity. Examples of relevant environmental risks known to be associated with ADHD are pregnancy and delivery complications, low birth weight and maternal alcohol consumption during pregnancy (51). In our model, these events lead to a cascade of events that cause: a) stressed neurons and astrocytes; b) dysregulated autophagy and apoptosis pathways and c) cell death. SBRV occurs because some brain regions are better protected from this cascade due to the degree to which they express genes in pathways that regulate apoptosis and autophagy. Those regions that are weakly protected suffer more cell loss than other regions leading to SBRV. Some cases of ADHD and ADHD associated volumetric reductions persist into adulthood (2, 3, 24, 52). Our results from the adult ADHD analyses suggest that the association of the neurodevelopmental gene set with SBRV in adult ADHD may account for this persistence. Because the neurodevelopmental gene set is involved in neurogenesis in adolescence and adulthood, we postulate that those brain regions that express these pathways to a greater degree are more likely than other brain regions to recover from SBRV.

A limitation of our study is that we provided only indirect evidence of pathways potentially relevant to SBRV in ADHD. No strong causal inferences can be drawn from the cross-sectional data used in our study. Ideally, studies of gene expression in brain tissue from ADHD patients could be used to confirm our results but no such tissue resources exist at this time.

Despite these limitations, our work is beginning to clarify potential causes of SBRV in ADHD and to provide hypotheses for testing in future research. By identifying specific cell types and biological pathways affected in ADHD, we provide guidance for future *in vitro* studies to test our theoretical model. Such studies may uncover genetic or environmental factors that influence SBRV and lead to cellular models that can be used to develop new medications for the disorder.

## Acknowledgements

SJG is supported by grants from the U.S. National Institutes of Health (5R01MH101519, 5R01AG054002), the Sidney R. Baer, Jr. Foundation, and NARSAD: The Brain & Behavior Research Foundation. This project has received funding from the European Union’s Horizon 2020 research and innovation programme grant agreement No 667302 (SVF).

