## Supplementary Tables for "Spatial organization of cells and variable expression of autophagy, apoptosis, and neurodevelopmental genes might underlie selective brain region vulnerability in Attention-Deficit/Hyperactivity Disorder"

**Affiliations:**

### Supplementary Tables

Supplementary Table 1. Gene set association analysis with data from the largest available GWAS meta-analysis of ADHD published by Demontis *et al*., (2018).

| **Source of gene set** | **Gene set** | **# of genes** | **Beta** | **SE** | ***p*-value** |
| --- | --- | --- | --- | --- | --- |
| Gene Ontology | Apoptosis | 35 | -0.05 | 0.16 | 0.615 |
|  | Autophagy | 353 | 0.03 | 0.05 | 0.237 |
|  | Neurodevelopment | 243 | 0.04 | 0.06 | 0.254 |
|  | Neurotransmission | 335 | 0.06 | 0.05 | 0.122 |
|  | Oxidative stress | 302 | 0.01 | 0.05 | 0.417 |
| Embryonic ventral midbrain scRNA-seq | DA0 | 38 | 0.20 | 0.15 | 0.096 |
|  | DA1 | 37 | 0.08 | 0.16 | 0.314 |
|  | DA2 | 49 | 0.05 | 0.13 | 0.337 |
|  | Endo | 60 | 0.11 | 0.13 | 0.184 |
|  | **GABA** | **61** | **0.23** | **0.12** | **0.029** |
|  | Mgl | 114 | 0.00 | 0.08 | 0.490 |
|  | NProg | 19 | -0.06 | 0.24 | 0.606 |
|  | NbGaba | 103 | -0.04 | 0.10 | 0.680 |
|  | NbM | 19 | -0.05 | 0.20 | 0.594 |
|  | NbML1 | 25 | 0.02 | 0.18 | 0.460 |
|  | NbML5 | 42 | -0.03 | 0.14 | 0.570 |
|  | OMTN | 14 | -0.05 | 0.30 | 0.561 |
|  | OPC | 70 | 0.03 | 0.12 | 0.393 |
|  | Peric | 39 | -0.04 | 0.16 | 0.599 |
|  | ProgBP | 19 | 0.06 | 0.21 | 0.388 |
|  | ProgFPL | 25 | 0.12 | 0.21 | 0.280 |
|  | ProgFPM | 32 | -0.02 | 0.17 | 0.545 |
|  | ProgM | 23 | -0.26 | 0.20 | 0.899 |
|  | RN | 42 | 0.08 | 0.14 | 0.288 |
|  | Rgl1 | 32 | -0.18 | 0.16 | 0.867 |
|  | Rgl2a | 15 | -0.15 | 0.22 | 0.750 |
|  | Rgl2b | 22 | 0.08 | 0.23 | 0.368 |
|  | Rgl2c | 45 | 0.13 | 0.14 | 0.162 |
|  | Rgl3 | 49 | 0.01 | 0.14 | 0.458 |
|  | Sert | 73 | -0.12 | 0.11 | 0.853 |
| Adult cortex scRNA-seq | OPC | 122 | 0.04 | 0.09 | 0.323 |
|  | astrocytes | 170 | -0.05 | 0.07 | 0.775 |
|  | endothelial | 166 | -0.09 | 0.08 | 0.887 |
|  | microglia | 180 | -0.01 | 0.07 | 0.545 |
|  | neurons | 184 | 0.04 | 0.07 | 0.272 |
|  | oligodendrocytes | 126 | -0.11 | 0.08 | 0.914 |

### Supplementary Figures

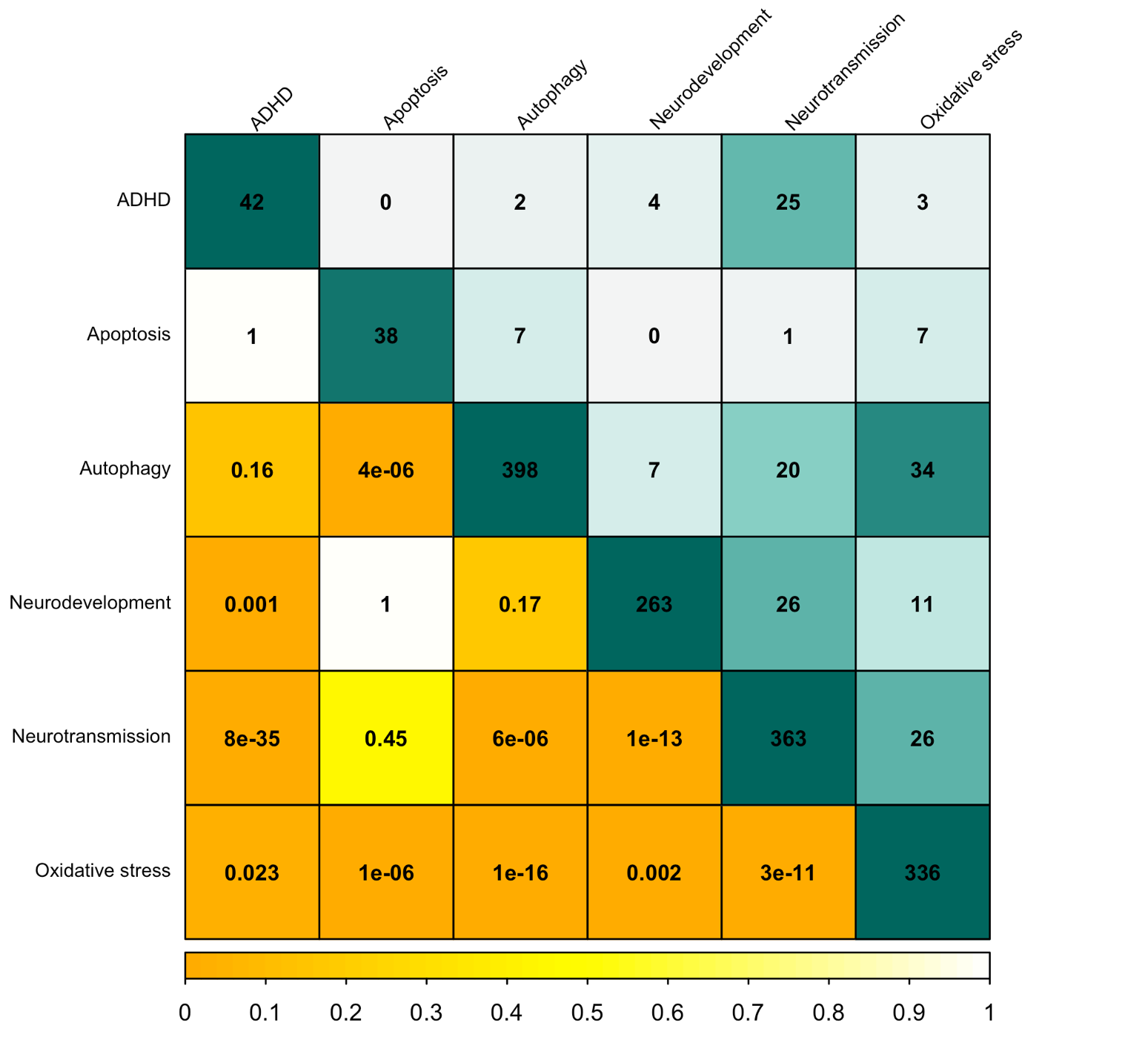

Supplementary Figure 1. The total number of genes within each gene set (along diagonal) along with the number of shared genes identified between pairs of gene sets (above diagonal). One-tailed Fisher’s *p*-values are provided below the diagonal, indicating whether the number of shared genes identified between pairs of gene sets was greater than expected by chance.

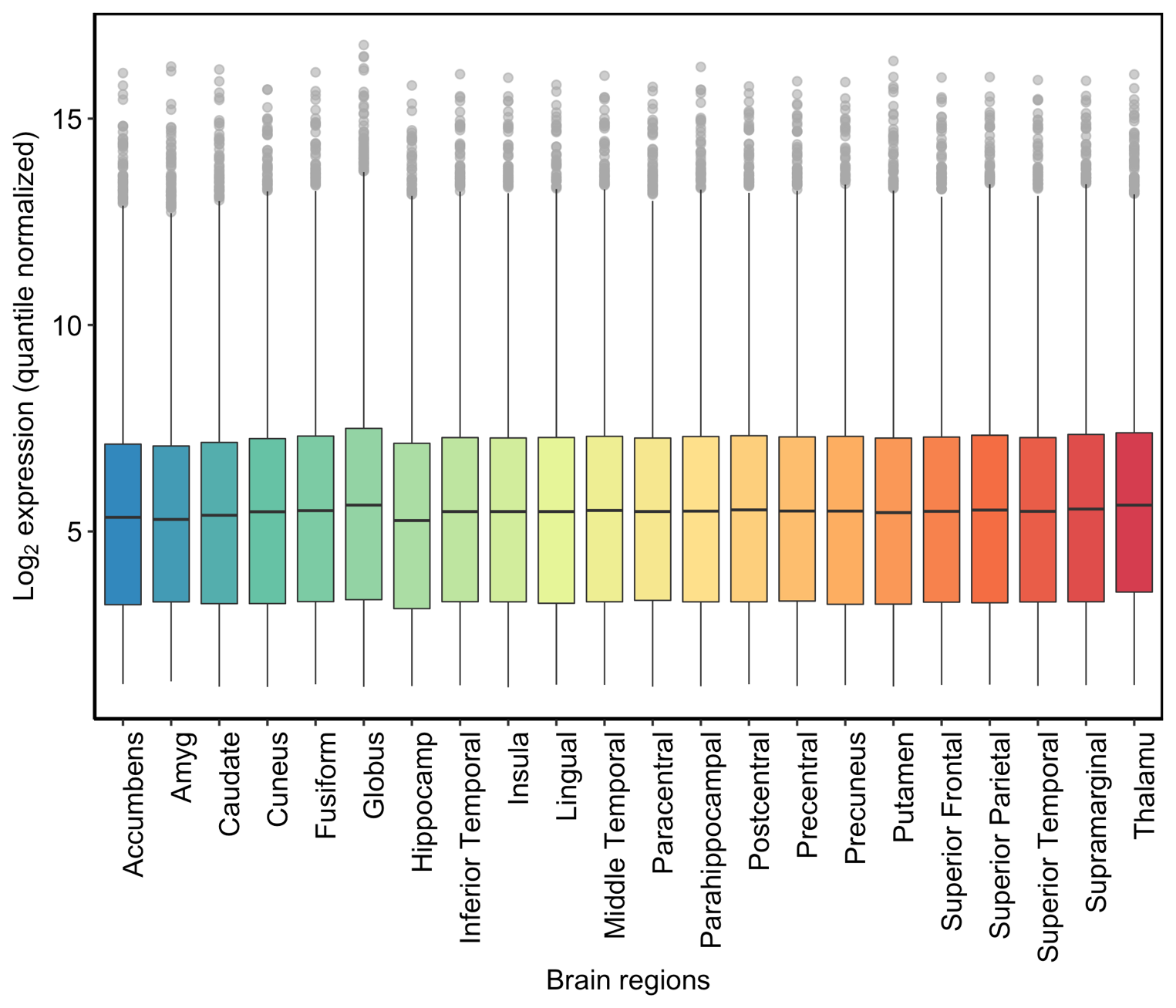

Supplementary Figure 2. Log_2_ quantile-normalized transcriptome profiles of 22 brain structures using data obtained from the Allen Brain Atlas (Shen et al., 2012).

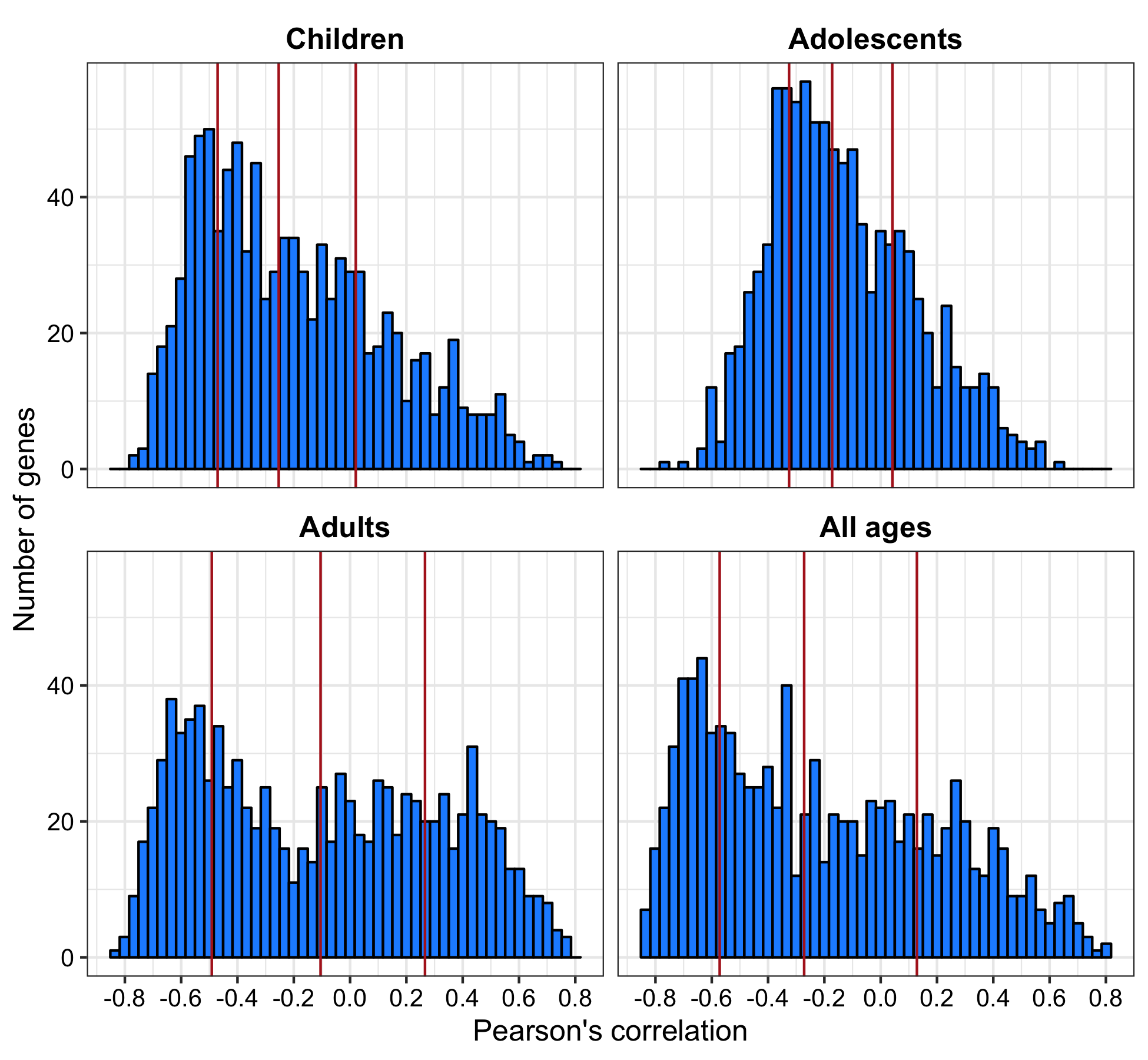

Supplementary Figure 3. Distribution of Pearson’s correlation coefficients (*r*) for 975 genes with brain volumetric changes in 22 sub-cortical and cortical brain regions in persons diagnosed with ADHD. Three red vertical lines denote the 25^th^, 50^th^, and 75^th^ percentile of Pearson’s correlations, in order, per age group.

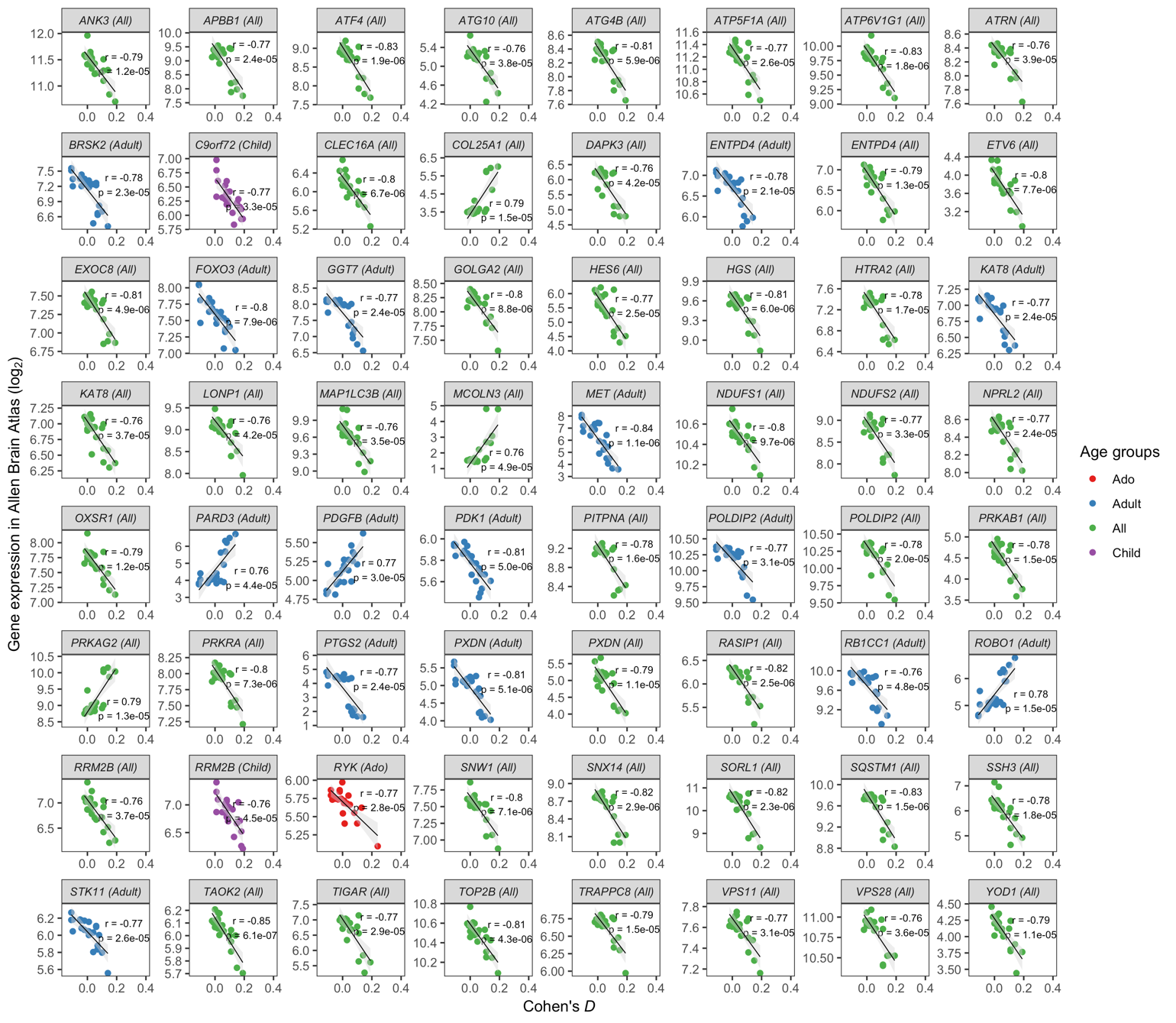

Supplementary Figure 4. Scatterplots depicting the significant correlations between gene expression levels for 64 genes and volumetric reductions ADHD cases. These associations had a Bonferroni-adjusted *p* < 0.05 after correcting for 974 genes tested within each of the four case-control age groups evaluated by the ENIGMA-ADHD working group. A best-fit regression line and shaded band denoting the standard error of the correlation is provided in each panel.

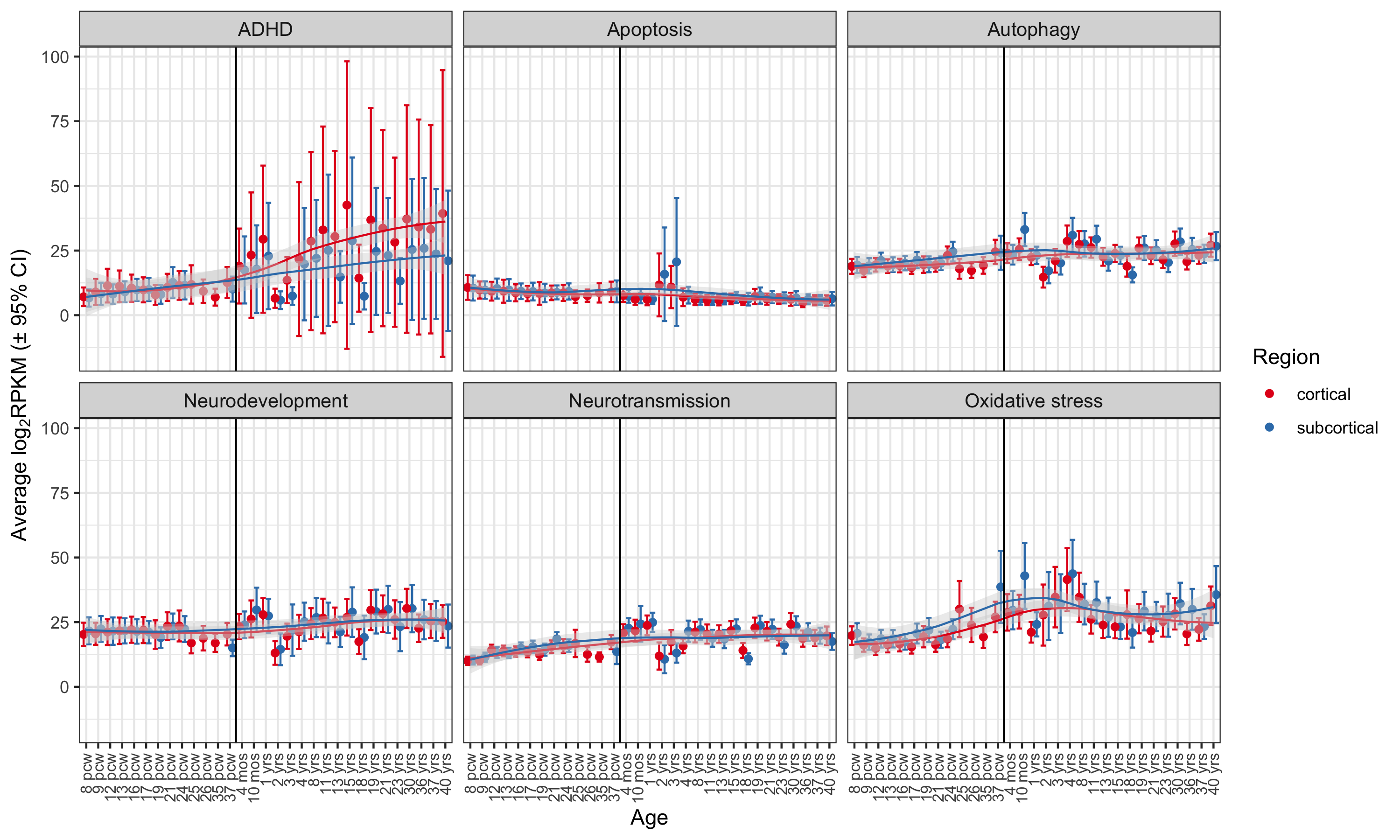

Supplementary Figure 5. Developmental trajectory of six gene sets in cortical and sub-cortical brain structures, with the average gene set expression level shown per age group. A vertical black line separates prenatal and postnatal periods. Red and blue lines represent LOESS regression fits per brain region. **Abbreviations:** Post-conceptual weeks (pcw), months (mos), years (yrs)
